# Prdm16 and Vcam1 regulate the postnatal disappearance of embryonic radial glia and the ending of cortical neurogenesis

**DOI:** 10.1101/2023.02.14.528567

**Authors:** Jiwen Li, Marlesa I. Godoy, Alice J. Zhang, Graciel Diamante, In Sook Ahn, Arantxa Cebrian-Silla, Arturo Alvarez-Buylla, Xia Yang, Bennett G. Novitch, Ye Zhang

## Abstract

Embryonic neural stem cells (NSCs, *i.e*., radial glia) in the ventricular-subventricular zone (V-SVZ) generate the majority of neurons and glia in the forebrain. Postnatally, embryonic radial glia disappear and a subpopulation of radial glia transition into adult NSCs. As this transition occurs, widespread neurogenesis in brain regions such as the cerebral cortex ends. The mechanisms that regulate the postnatal disappearance of radial glia and the ending of embryonic neurogenesis remain poorly understood. Here, we show that PR domain-containing 16 (Prdm16) promotes the disappearance of radial glia and the ending of neurogenesis in the cerebral cortex. Genetic deletion of *Prdm16* from NSCs leads to the persistence of radial glia in the adult V-SVZ and prolonged postnatal cortical neurogenesis. Mechanistically, Prdm16 induces the postnatal reduction in Vascular Cell Adhesion Molecule 1 (Vcam1). The postnatal disappearance of radial glia and the ending of cortical neurogenesis occur normally in *Prdm16-Vcam1* double conditional knockout mice. These observations reveal novel molecular regulators of the postnatal disappearance of radial glia and the ending of embryonic neurogenesis, filling a key knowledge gap in NSC biology.

## Introduction

The mammalian brain has limited regenerative capacity compared with fish, amphibian, and reptile brains^1^. In the adult mouse brain, new neurons are generated in only two brain regions: the olfactory bulb and the dentate gyrus^2,3^. In humans, the existence and extent of adult neurogenesis remains under debate^4,5^. Stroke, traumatic brain injury, and neurodegenerative disorders lead to neuronal death. In these cases, the growth of new axon collaterals from surviving neurons and the reorganization of neural circuits can help patients regain some lost function. However, few new neurons are generated to rebuild damaged neural circuits^6^. Therefore, most patients suffer from permanent motor and/or cognitive dysfunction, which reduces quality of life. In contrast to the limited neurogenesis capacity in adults, the mammalian brain is highly efficient in generating neurons during embryonic development. For example, billions of new neurons are generated within months in the human fetal brain. Embryonic neural stem cells (NSCs, *i.e*., radial glia) are responsible for rapid and widespread embryonic neurogenesis^7^. The mechanisms regulating the postnatal disappearance of radial glia and the ending of brain-wide neurogenesis remain elusive.

Radial glia reside in the V-SVZ on the wall of the lateral ventricle and generate the majority of neurons and glia in the forebrain^8–14^. Each NSC/radial glia has a long radial process that frequently extends deep into the adjacent brain parenchyma. Many of these radial glia, and especially those in the cortical walls have endfeet at the pial surface^13–17^. In the cortex, radial processes guide the migration of new neurons generated in the V-SVZ to their final destination in the cortical plate^7,10,14,16–23^.

From embryonic day 13.5 to 15.5 (E13.5-15.5), a subpopulation of embryonic NSCs/radial glia slows down cell division and are “set aside” to become adult NSCs (type B cells) responsible for adult neurogenesis in mouse^24–27^. The rest of radial glia continue to divide rapidly and contribute to embryonic neurogenesis. We refer to the subpopulation of embryonic radial glia destined to become adult type B cells as “slowly dividing” radial glia and other radial glia that generate embryonic neurons and glia as “rapidly dividing” radial glia. In the first postnatal month, rapidly dividing radial glia differentiate and disappear, whereas slowly dividing radial glia disappear by transitioning into adult NSCs^24–26,28^. During the postnatal transition, the long radial processes retract. The adult NSCs/B cells derived from slowly dividing radial glia are generally multipolar with a short basal process (a process oriented towards the brain parenchyma) that ends in neighboring blood vessels^29^. Unlike embryonic radial glia, which guide the dorsal migration of cortical excitatory neurons with their radial processes, adult B cells lack radial processes and do not guide neuronal migration. As radial glia transition into adult NSCs, the types of neurons they generate also change. V-SVZ radial glia generate cortical excitatory neurons and striatal neurons, whereas adult B cells generate olfactory bulb interneurons^10,11,30^ Neurogenesis in most brain regions (except the olfactory bulb and dentate gyrus) ends postnatally, as radial glia transition into adult B cells. Similarly, radial glia in humans (outer radial glia and ventricular radial glia) are responsible for the rapid generation of neurons and the guidance of neuronal migration during embryonic development^15,31^. Postnatally, radial glia with long radial processes disappear, and neurogenesis in brain regions such as the cerebral cortex ends^32,33^. Although there is an ongoing debate on whether and to what extent adult hippocampal neurogenesis occurs in humans similar to mice^4,5^, both species exhibit the postnatal disappearance of radial glia and the ending of brain-wide neurogenesis. Thus, uncovering the mechanisms that regulate postnatal changes of NSCs in mice will provide a foundation for future studies that improve our understanding of human NSC biology and NSC-based therapies.

Although the postnatal disappearance of radial glia and the ending of neurogenesis in most brain regions have been observed for decades, the molecular signals that trigger these postnatal changes remain poorly understood. Recent studies have provided an increasingly detailed understanding of the regulation of embryonic neurogenesis and adult neurogenesis^10,34^. The postnatal transition process, however, remains one of the least, if not *the* least, understood stage in the lifespan of NSCs.

Serendipitous discoveries led us to investigate the potential role of PR-domain containing 16 (Prdm16) in the transition from embryonic to adult NSCs. Prdm16 is an epigenetic regulator with histone methyltransferase activity^35,36^. Studies have shown that Prdm16 is important for the maintenance of neural and hematopoietic stem cells^37,38^ and for the development of upper-layer cortical neurons^39–41^ interneurons^42^, and vascular cells^43^. These studies focused on the embryonic^37,39–41,43^ and adult stages^38^ and the function of Prdm16 during the early postnatal stage in NSCs has not been reported. Whether Prdm16 plays a role in the postnatal disappearance of radial glia and the ending of embryonic neurogenesis remains unknown.

Here, we found that radial glia with long basal processes persist in adulthood and postnatal cortical neurogenesis is prolonged in *Prdm16-conditional* knockout (cKO) mice compared with controls, demonstrating that Prdm16 is indispensable for the postnatal disappearance of radial glia and the ending of cortical neurogenesis. Furthermore, we found that Prdm16 induces a postnatal reduction in the level of Vascular Cell Adhesion Molecule 1 (Vcam1) and that inhibition of Vcam1 by Prdm16 is essential for the postnatal transition. Hence, we identified Prdm16 and Vcam1 as key regulators for the postnatal disappearance of radial glia and the ending of cortical neurogenesis.

## Results

### The basal processes of embryonic NSCs/radial glia persist in adulthood in *Prdm16-cKO* mice

We first assessed the distribution of Prdm16 protein in the postnatal brain and detected it in the V-SVZ (Fig. 1A, postnatal day 16), consistent with previous reports^37,39^. To investigate the potential roles of Prdm16 in the postnatal transition from embryonic to adult NSCs, we crossed *human-GFAP-Cre (hGFAP-Cre*) transgenic mice, which express Cre recombinase in radial glia starting from embryonic day 13.5 (E13.5)^44^ - the time at which a subpopulation of radial glia slow down cell division and are set aside to become adult B cells for adult neurogenesis - with *Prdm16*-floxed knock-in mice^45^ to generate *hGFAP-Cre^T9/+^; Prdm16^fl/fl^* cKO mice. *Prdm16* cKO mice are viable and fertile. We used *Prdm16^fl/fl^* littermates without *hGFAP-Cre* as controls. We detected reduced levels of Prdm16 protein in the postnatal V-SVZ of cKO mice (Fig. 1A).

**Figure 1.**
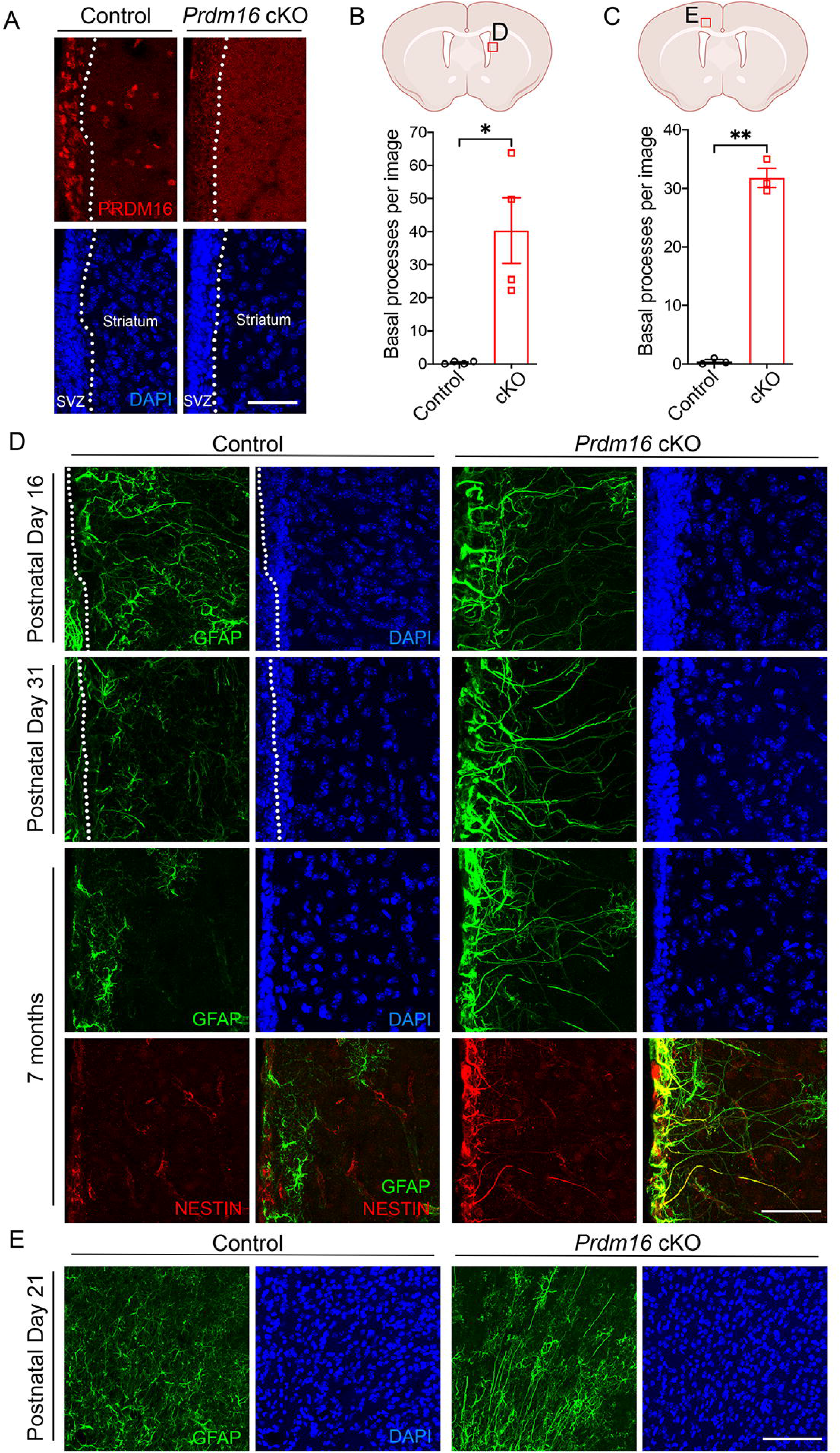
Persistence of cells with radial processes in juvenile and adult *Prdm16* cKO mice. (A) Prdm16 immunofluorescence in the V-SVZ and striatum of *Prdm16* cKO and control mice at P16. The dashed lines delineate the boundaries between the V-SVZ and striatum. Scale bar: 50 μm. (B) A diagram shows the area of lateral V-SVZ used in the quantification and shown in (D). Quantification of the number of radial processes at one month of age in the striatum. Images were taken between −0.1 mm posterior and 0.74 mm anterior to Bregma. Only radial processes that extend beyond 200 μm away from the ventricular walls were counted. N=4 images per mouse, 4 mice per genotype. All statistical tests were performed using data from each mouse (biological replicate) as an individual observation in all figures except for scRNAseq. Welch’s t-test. p=0.0276. In all figures: *, p<0.05. **, p<0.01. ***, p<0.001. N.S., not significant. (C) A diagram shows the area of the cerebral cortex used in the quantification and shown in (E). Quantification of the number of radial processes at P21 in the cerebral cortex. Images were taken between 0.1 and 1.18 mm anterior to Bregma. Only radial processes that extend beyond the white matter were counted. N=5-8 images per mouse, 3 mice per genotype. Welch’s t-test. p=0.0021. (D) Persistence of GFAP^+^/Nestin^+^ basal processes in juvenile and adult *Prdm16* cKO mice. Scale bar: 50 μm. The dashed lines indicate ventricular walls. (E) Persistence of GFAP^+^ basal processes in P21 *Prdm16* cKO cortex. Scale bar: 100 μm.

We next assessed the presence of embryonic NSC/radial glia in postnatal *Prdm16* cKO and control mouse brains by immunostaining against GFAP and Nestin, markers of radial glia. In control mice, basal processes of radial glia are abundant in neonates but mostly retract by P16. Strikingly, basal processes that extend beyond 200 μm away from the ventricular wall persist in juvenile (one-month-old) and adult (seven-month-old) *Prdm16* cKO mice (Fig. 1B-E). We observed similar persistence of radial processes in the striatum (Fig. 1B, D) and the cerebral cortex (Fig. 1C, E). This interesting observation demonstrates that Prdm16 is required for the postnatal retraction of the radial processes of radial glia.

### NSCs are increased and ependymal cells are decreased in the V-SVZ of *Prdm16* cKO mice

Two major cell types, type B cells and ependymal cells are located on the surface of the walls of the lateral ventricles in adult mice^52^. Type B cells are adult NSCs^2^, whereas ependymal cells produce signals that regulate NSC proliferation and have multiple cilia that propel the circulation of cerebrospinal fluid in the ventricles^53,54^. NSCs are GFAP^+^ and Vcam1^+^ and contain a primary cilium associated with basal bodies (labeled by γ-tubulin)^55^. Ependymal cells are Vcam1^-^ and contain multiple *γ*-tubulin^+^ basal bodies connected to an array of motile cilia (Fig. 2A, B)^54^. Adult type B cells and ependymal cells are produced by shared embryonic radial glia progenitors^26,27^. To examine the cell composition of the surface of the lateral wall of the lateral ventricle, we made whole-mount preparations which expose the surface of the lateral wall at one month of age *Prdm16* cKO and control mice and stained them with antibodies against Vcam1, γ-tubulin, and Sox2 (which labels the nuclei of both NSCs and ependymal cells and is helpful for cell counting) (Fig. 2C). Interestingly, *Prdm16* cKO mice showed an increase in NSCs and a decrease in ependymal cells (Fig. 2D, E). In addition, we found that *Prdm16* cKO mice often exhibit hydrocephalus, a phenotype commonly observed in mice with ependymal cell defects^56,57^ (Supplementary Fig. 1). The increase in NSCs in *Prdm16* cKO mice is a novel finding and is consistent with our hypothesis that radial glia persist postnatally in these mice. The decrease in ependymal cells replicates a prior report^38^.

**Figure 2.**
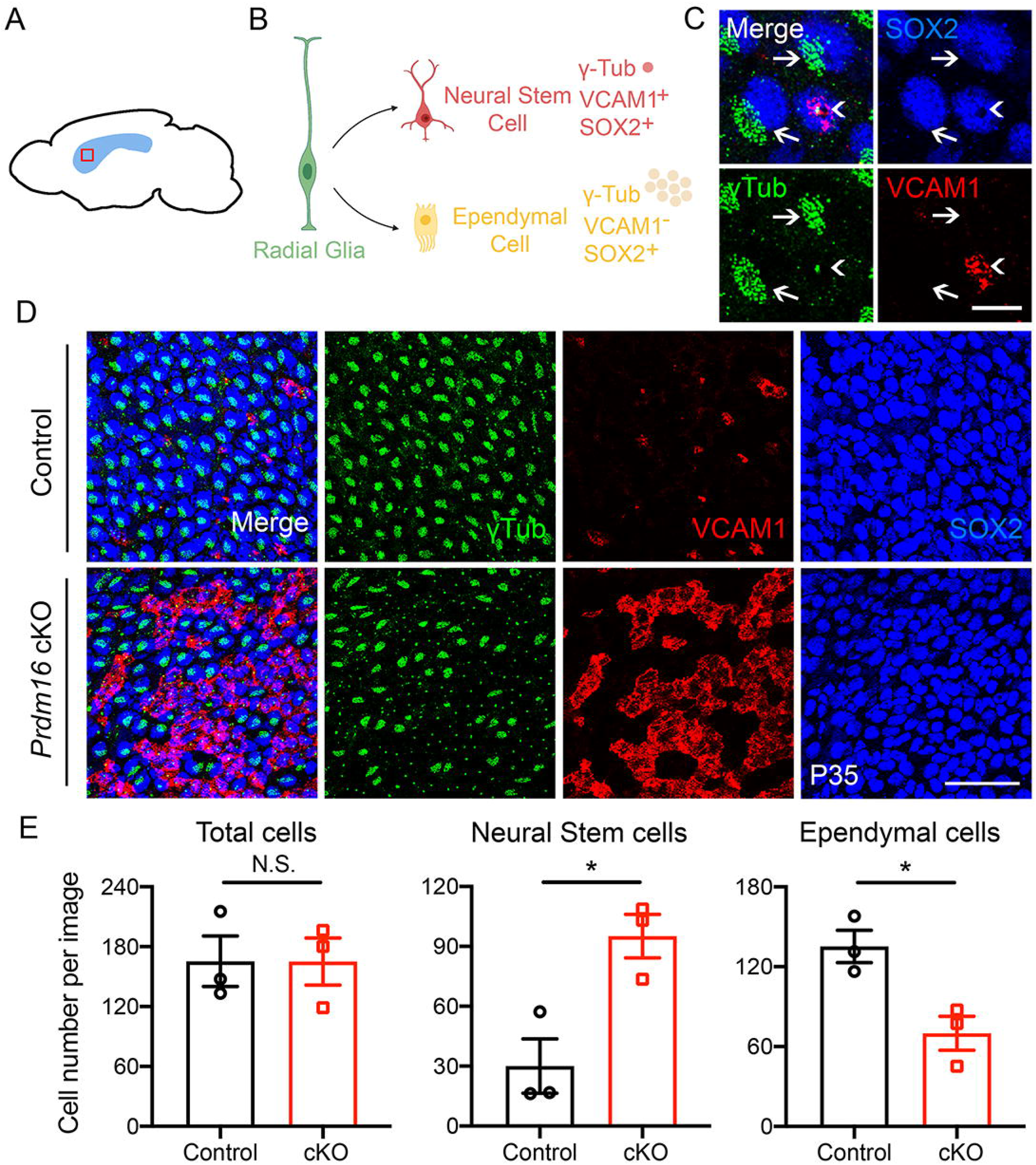
An increase in neural stem cells and a decrease in ependymal cells in *Prdm16* cKO V-SVZ. (A) Whole-mount preparation, in which the entire surface of the lateral walls of the lateral ventricles is cut out for immunohistochemistry experiments. (B) Neural stem cells and ependymal cells were identified by a combination of markers on whole-mount preparations. Neural stem cells are positive for Vcam1 and Sox2 with single dots positive for γ-tubulin immunofluorescence (a single basal body associated with a single primary cilium). Ependymal cells are negative for Vcam1 and GFAP with clusters of γ-tubulin immunofluorescence (multiple basal bodies associated with multiple cilia). (C) An example of a Vcam1^+^ neural stem cell with a single γ-tubulin^+^ dot (arrowhead) and examples of Vcam1^-^ ependymal cells with clusters of γ-tubulin^+^ dots (arrows). Scale bar: 10 μm. (D) Vcam1, γ-tubulin, and Sox2 immunofluorescence on whole-mount preparations of P35 *Prdm16*-knockout and control mice. Scale bar: 50 μm. (E) Quantification of neural stem and ependymal cells. N=4 images per mouse, 3 mice per genotype. Welch’s t-test, neural stem cells p=0.0219, ependymal cells p=0.0208.

### Bromodeoxyuridine (BrdU) pulse chase experiments reveal the postnatal persistence of slowly dividing embryonic radial glia in *Prdm16* cKO mice

The subpopulation of radial glia set aside to become adult type B cells divide slowly whereas other embryonic radial glia divide rapidly ^24,25^. A previous study demonstrated that BrdU labelling is quickly diluted with each cell cycle in rapidly dividing radial glia and the only cells retaining high levels of BrdU several weeks after BrdU injection are slowly dividing radial glia ^46^. To directly trace the postnatal fate of slowly dividing embryonic radial glia, we performed BrdU pulse-chase experiments by injecting BrdU between E15.5 and E17.5 and sacrificing the mice at P21 (Fig. 3A). Interestingly, the number of BrdU-labeled cells in the V-SVZ was higher in *Prdm16* cKO mice compared with control mice at P21 (Fig. 3B-C), lending further support to our hypothesis that slowly dividing embryonic radial glia persist postnatally in *Prdm16* cKO mice.

**Figure 3.**
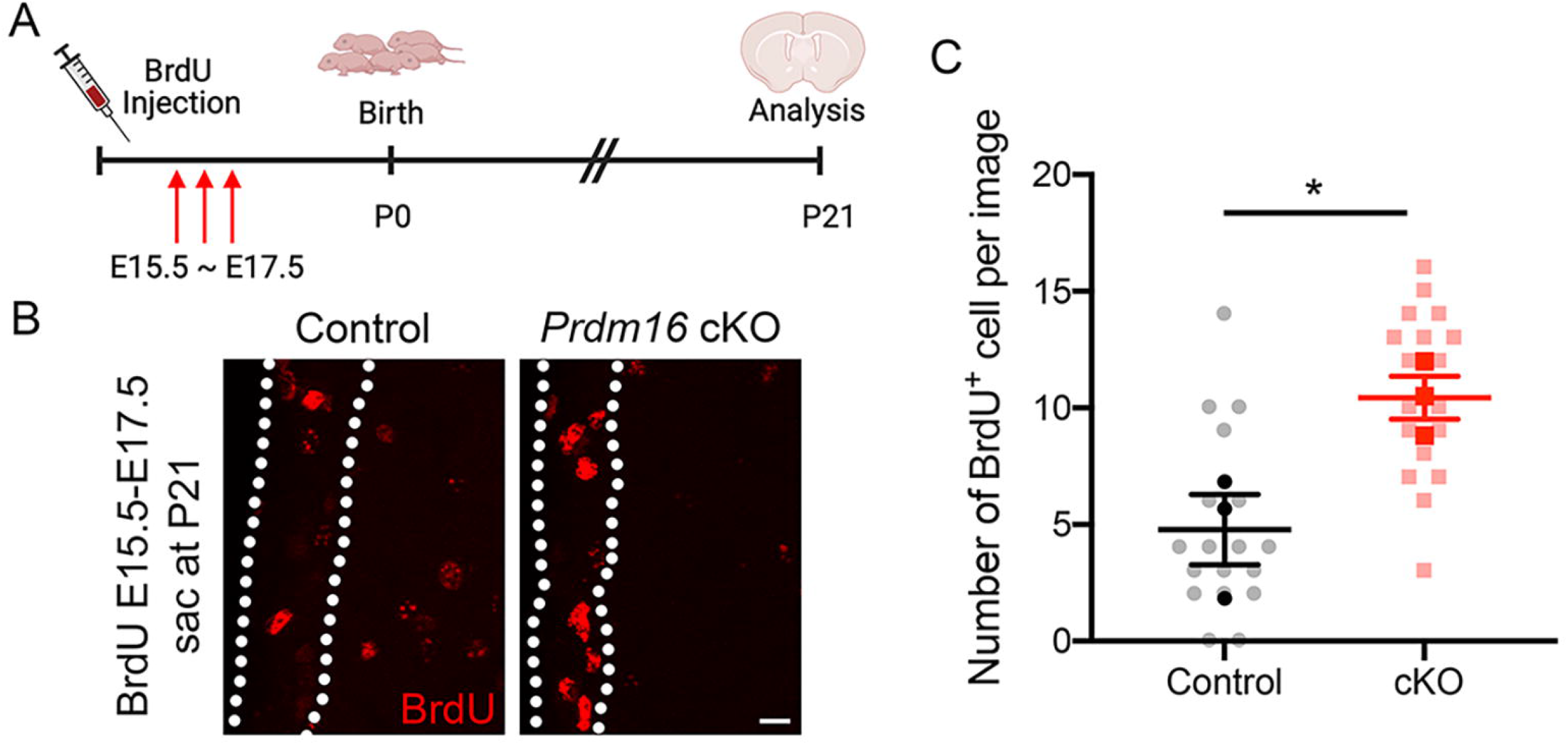
Embryonic slowly dividing radial glia persist postnatally in the V-SVZs of *Prdm16* cKO mice. (A) A diagram shows the design of the BrdU pulse-chase experiment. BrdU was injected 3 times daily between E15.5 and E17.5, and the mice were sacrificed and analyzed at P21. (B) Increased BrdU-labeled cells in the V-SVZs of *Prdm16* cKO mice compared with control at P21. The dashed lines delineate V-SVZs. Scale bar: 10 μm. (C) The quantification of BrdU-labeled cells in the V-SVZ. N=5-7 images per mouse, 3 mice per genotype. The black and red dots indicate average results from each mouse and the grey and pink dots indicate the raw counts from each image. Welch’s t-test, p=0.0431.

### Single-cell RNA sequencing (scRNAseq) analysis reveals postnatal persistence of slowly dividing radial glia in Prdm16 cKO mice

To comprehensively assess the molecular changes in the V-SVZ of *Prdm16* cKO mice, we performed scRNAseq from one-month-old V-SVZ. We micro-dissected the lateral walls of the lateral ventricles using the same method described for the preparation of wholemounts^47^, dissociated the tissue, and performed scRNAseq with the 10x Genomics platform using cells from three cKO and three control mice. After filtering out low-quality cells, we obtained transcriptome data for 36,420 V-SVZ cells, including 21,939 and 14,481 cells from control and *Prdm16* cKO mice, respectively. We next performed unbiased cell clustering and used uniform manifold approximation and projection (UMAP) for data visualization. We annotated the identities of cell clusters based on known cell-type-specific markers (Fig. 4A, B). The sequenced V-SVZ cells included all expected cell types identified in published scRNAseq datasets of the V-SVZ^48–51^, such as NSCs, type C cells/transiently amplifying precursors (TAPs), type A cells/neuroblasts, neurons, ependymal cells, astrocytes, oligodendrocyte precursor cells, oligodendrocytes, microglia, endothelial cells, and pericytes (Fig. 4A, B, and Supplementary Fig. 2A-C). When we compared *Prdm16* cKO vs control mice, we detected differentially expressed genes in all cell types, with NSCs exhibiting the most robust transcriptome changes (Fig. 4C and Tables S1, S2).

**Figure 4.**
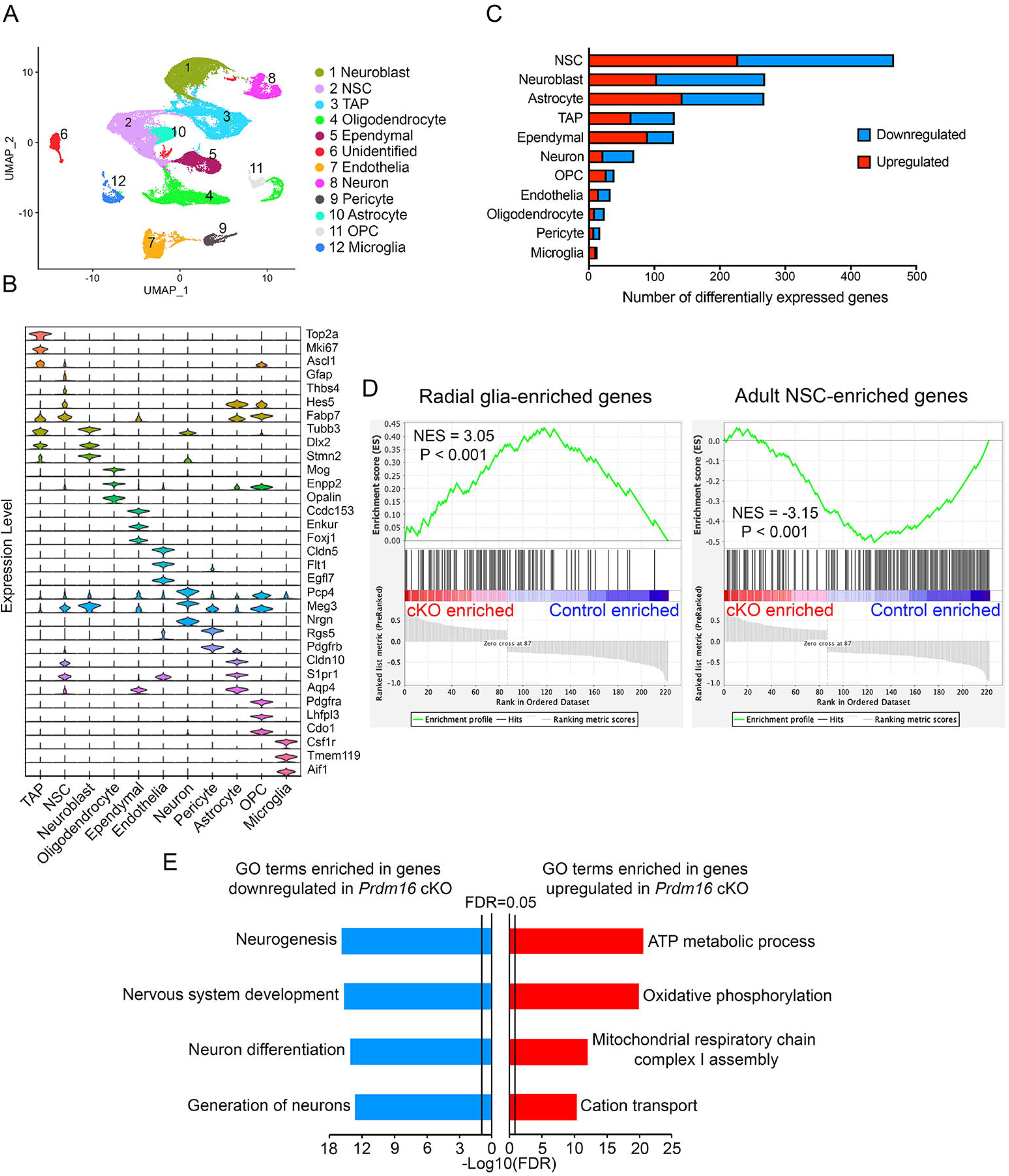
ScRNAseq analysis of the V-SVZs of *Prdm16* cKO mice. (A) UMAP of scRNAseq data. Major cell clusters were identified and color-coded. (B) Expression of cell-type markers by the major cell types. (C) The numbers of differentially expressed genes in each cell type. (D) Prdm16 cKO NSCs upregulated the expression of embryonic radial glia-enriched genes and downregulated the expression of adult NSCs-enriched genes. NES: normalized enrichment score. (E) Selected GO terms enriched in genes up- and down-regulated in *Prdm16* cKO NSCs compared with controls

We next assessed whether NSCs in *Prdm16* cKO mice display a transcriptome-wide delay in the transition from the slowly dividing embryonic radial glia to the adult B cells. Based on the literature, slowly dividing radial glia and adult B cells share the expression of NSC marker genes. Therefore, we performed an analysis to identify gene expression patterns that may help us distinguish between slowly dividing radial glia and adult B cells. We used a published embryonic and adult brain scRNAseq dataset^48^. We first extracted the E14 embryonic radial glia cluster data and generated subclusters (Supplementary Fig. 3 and Supplementary Fig. 4A). Based on proliferation marker genes, we identified a slowly dividing radial glia subcluster and a rapidly dividing radial glia subcluster (Supplementary Fig 4B-D). We then compared the gene expression of slowly dividing radial glia *vs*. adult type B cells and identified differentially expressed genes (RGvB-DEGs). To assess the expression of RGvB-DEGs in *Prdm16* cKO and control NSCs, we next performed gene set enrichment analysis (GSEA). We used genes with higher expression in radial glia than adult B cells as one gene set and genes with lower expression in radial glia than adult B cells as a second gene set.

Interestingly, we found that the genes upregulated in *Prdm16* cKO NSCs are enriched in genes expressed at higher levels in radial glia than adult type B cells whereas the genes downregulated in Prdm16 *cKO* NSCs are enriched in genes expressed at lower levels in radial glia than adult type B cells (Fig. 4D, p<0.001), suggesting a delay in the transition of the transcriptome profile from embryonic radial glia to adult type B cells.

Together, the three lines of evidence described above suggest that the transition from embryonic to adult NSCs is delayed in *Prdm16* cKO mice: (1) cells with the morphology of embryonic radial glia (long radial processes) persist in adulthood (Fig. 1), (2) slowly dividing cells labeled with BrdU at E15.5-17.5 are increased in the V-SVZs of juvenile *Prdm16* cKO mice (Fig. 3), and (3) the transcriptome profile of juvenile *Prdm16* cKO NSCs resembles that of embryonic radial glia (Fig. 4).

### The expression of oxidative phosphorylation-associated genes is increased in *Prdm16* cKO NSCs

To gain insight into the function of genes differentially expressed in *Prdm16* cKO mice, we performed gene ontology (GO) and protein-protein interaction network analyses. Among the upregulated genes, 23 out of 228 are within a protein-protein interaction network involved in oxidative phosphorylation (Supplementary Fig. 5), and the GO term oxidative phosphorylation is significantly enriched among the upregulated genes (Fig. 4E and Table S3). These genes encode mitochondrial electron transport chain components among others (e.g. *Ndufa9, Uqcr10, and Uqcrb*), suggesting that Prdm16 inhibits the expression of genes involved in oxidative phosphorylation. Our results uncover a potential role of Prdm16 in regulating energy metabolism in NSCs. GO terms enriched in genes downregulated in Prdm16 cKO NSCs include “nervous system development” and “neurogenesis” (Fig. 4E and and Table S3), consistent with a role of Prdm16 in NSC development. We also performed Gene Set Enrichment Analysis (GSEA) and obtained similar results as the GO term enrichment analysis (Supplementary Fig. 6).

### Cortical neurogenesis continues for longer postnatally in *Prdm16* cKO mice than in control mice

Embryonic radial glia generate cerebral cortical excitatory neurons and guide neuronal migration through the cortical layers with their long radial processes^2^. Shortly after birth, cortical neurogenesis ends ^58^, and newly generated neurons from adult B cells migrate to the olfactory bulb via the rostral migratory stream ^11,59^ The mechanisms that regulate the ending of cortical neurogenesis remain unidentified. Having observed the postnatal persistence of cells with long radial processes in Prdm16 cKO mice, we next examined whether these cells support the continued generation and migration of new cortical neurons by staining for doublecortin (DCX), a marker for newly generated neurons. At P21, few DCX^+^ cells were detected in the cortex of control mice, consistent with prior reports ^58^. Interestingly, we found that numerous DCX^+^ cells were present in the cerebral cortex of *Prdm16* cKO mice. These cells were most frequently seen in the secondary motor cortex and anterior cingulate cortex, the areas where persistent radial processes were most abundant (Fig. 5A-D). Some Dcx^+^ cells were present along GFAP^+^ radial processes, possibly migrating along them (Fig. 5C). These results suggest that Prdm16 promotes the ending of cortical neurogenesis.

**Figure 5.**
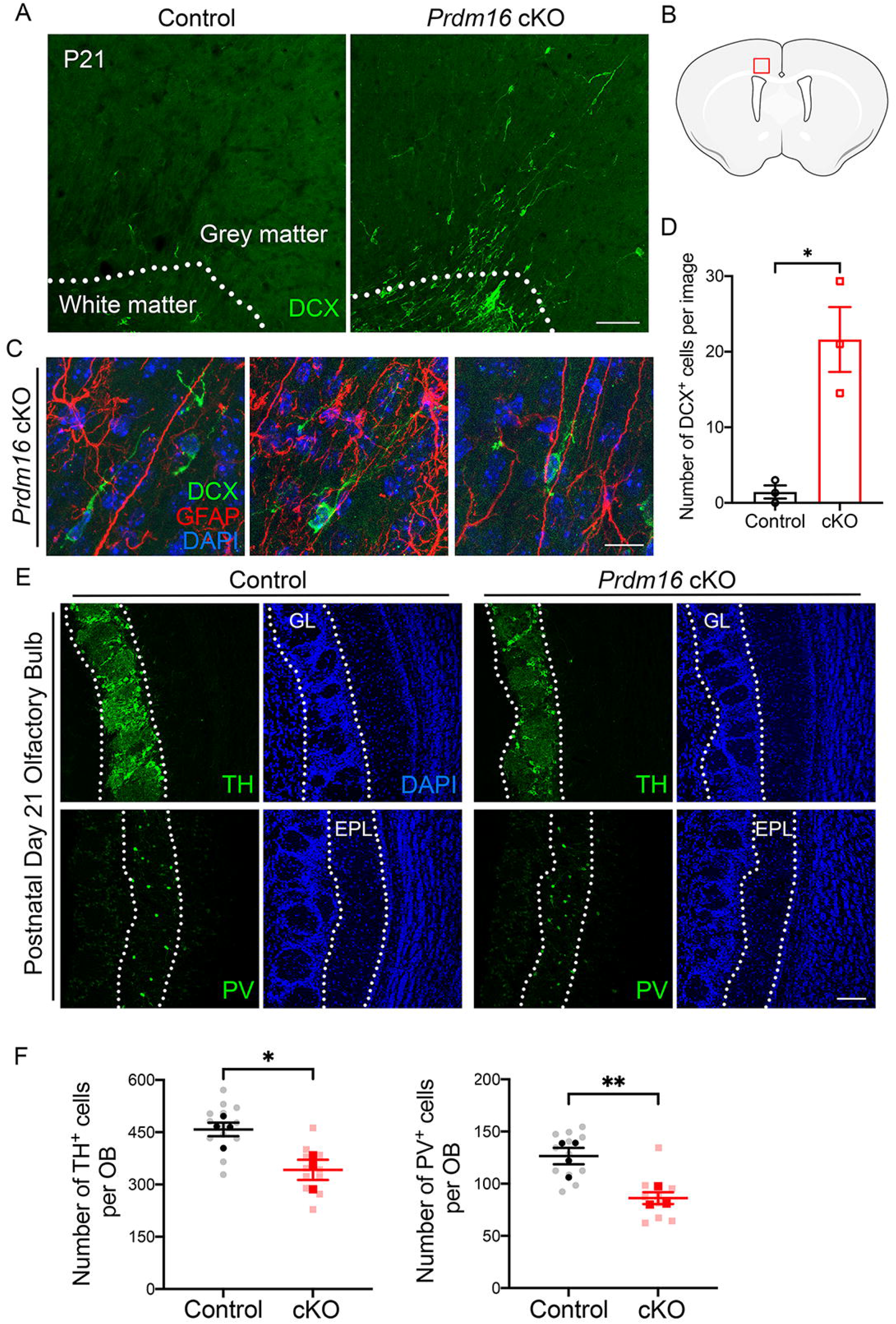
Postnatal persistence of cortical neurogenesis and olfactory bulb interneuron development defect in *Prdm16* cKO mice. (A) Numerous DCX^+^ newly born neurons in *Prdm16* cKO cerebral cortices at P21. Few new neurons are detected in control cerebral cortex at this age. The dashed lines delineate the boundaries between the white matter and the grey matter. Scale bar: 100 μm. (B) A diagram shows the area shown in (A) and (C) and quantified in (D). (C) DCX^+^ newborn neurons alongside GFAP^+^ radial processes in the cerebral cortex of *Prdm16* cKO mice at P21. Scale bar: 20 μm. (D) Quantification of the number of DCX^+^ cells at P21 in the cortex. Only DCX^+^ cells in the grey matter were counted. N=5-8 images per mouse, 3 mice per genotype. Welch’s t-test. p=0.0379. (E) Two subtypes of olfactory bulb interneurons. The densities of TH^+^ and PV^+^ interneurons were reduced in the olfactory bulbs of *Prdm16* cKO mice at P21. The dashed lines delineate the glomerular layer (GL), external plexiform layer (EPL), and granule cell layer (GCL) in the olfactory bulb. Scale bar: 100 μm. **(F)** Quantification of the two subtypes of olfactory bulb interneurons. N=1-5 images per mouse, 3-4 mice per genotype. The black and red dots indicate average results from each mouse and the grey and pink dots indicate the raw counts from each image. Welch’s t-test, TH^+^ cells, p=0.0137, PV^+^ cells, p=0.0092

### Tyrosine hydroxylase (TH)^+^ and parvalbumin (PV)^+^ olfactory bulb interneurons are reduced in Prdm16 cKO mice

To examine the development of olfactory bulb interneurons, we performed immunostaining at P21 with antibodies against markers of two subtypes of olfactory bulb interneurons: TH^+^ and PV^+^ interneurons. Interestingly, the densities of both TH^+^ and PV^+^ interneurons were reduced in *Prdm16* cKO mice (Fig. 5E-F), demonstrating a role of Prdm16 in the regulation of olfactory bulb interneuron development.

### Prdm16 promotes the postnatal disappearance of radial glia and the ending of cortical neurogenesis by repressing Vcam1

To investigate the mechanisms through which Prdm16 regulates the disappearance of embryonic radial glia, we took a candidate-based approach. We searched the literature for candidate genes/proteins exhibiting dynamic level changes during the transition period in NSCs. *Vcam1* is expressed by both embryonic and adult NSCs^46,60^. Interestingly, the Vcam1 protein is widespread in the embryonic germinal zone and the distribution becomes more restricted in the V-SVZ during the first three postnatal weeks as radial glial transition into adult NSCs^46^. Importantly, Vcam1 maintains the subpopulation of embryonic NSCs destined to become adult neural stem cells^46^; NSCs are prematurely depleted in *Vcam1*-knockout mice^60^. Furthermore, Vcam1 is necessary for maintaining the V-SVZ structure in adults^61^. However, it is unclear whether Vcam1 regulates the postnatal disappearance of radial glia or the ending of cortical neurogenesis.

To assess whether Prdm16 regulates Vcam1, we examined Vcam1 protein levels in the *Prdm16* cKO mice by immunohistochemistry. Interestingly, we found that juvenile and adult *Prdm16* cKO mice show higher levels of Vcam1 in the V-SVZ compared with controls (Fig. 6A, D).

**Figure 6.**
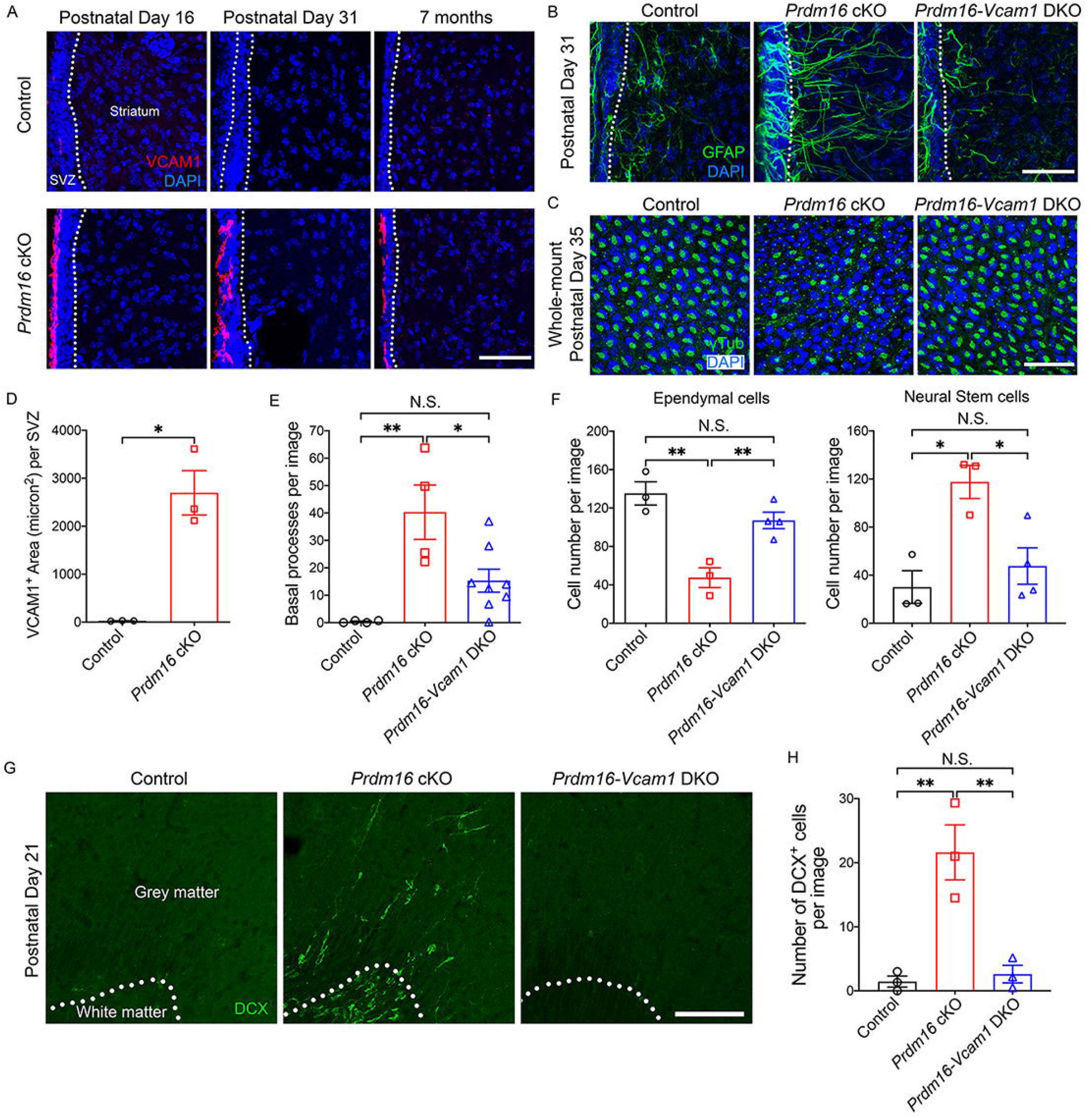
Rescue of the postnatal persistence of radial glia and cortical neurogenesis phenotypes in *Prdm16-Vcam1* DKO mice. (A) Increased Vcam1 immunofluorescence in the V-SVZ of juvenile and adult *Prdm16* cKO mice. The dashed lines delineate the boundaries between the V-SVZ and the striatum and the boundary between the V-SVZ and the ventricle. Scale bar: 50 μm. (B) The persistence of GFAP^+^ radial processes is partially rescued in *Prdm16-Vcam1* DKO mice. Scale bar: 50 μm. (C) Whole-mount en-face preparations showing neural stem cells with single γ-tubulin+ dots and ependymal cells with clusters of γ-tubulin+ dots. Scale bar: 50 μm. The cell composition defect in *Prdm16* cKO mice is rescued in *Prdm16-Vcam1* DKO. (D) Quantification of Vcam1 immunofluorescence positive area in V-SVZ. N=4-6 images per mouse, 3 mice per genotype. Welch’s t-test, p=0.0287. (E) Quantification of the number of radial processes at one month of age. N=4 images per mouse, 4-8 mice per genotype. One-way ANOVA with Tukey’s multiple comparisons test, control vs. *Prdm16* cKO, p=0.002, *Prdm16* cKO vs. *Prdm16-Vcam1* DKO, p=0.019. (F) Quantification of neural stem and ependymal cells. N=4-5 images per mouse, 3 mice per genotype. One-way ANOVA with Tukey’s multiple comparisons test. Ependymal cells, control vs. *Prdm16* cKO, p=0.0017, *Prdm16* cKO vs. *Prdm16-Vcam1* DKO, p=0.0097. Neural stem cells, control vs. *Prdm16* cKO, p=0.0124. *Prdm16* cKO vs. *Prdm16-Vcam1* DKO, p=0.0262. (G) The postnatal persistence of newly born DCX^+^ neurons in the cerebral cortex seen in *Prdm16* cKO mice is rescued in *Prdm16-Vcam1* DKO mice. The dashed lines delineate the boundaries between the white matter and the grey matter. Scale bar: 100 μm. (H) Quantification of DCX^+^ cells at P21 in the cerebral cortex. Only DCX^+^ cells in the grey matter were counted. N=5-8 images per mouse, 3 mice per genotype. Oneway ANOVA with Tukey’s multiple comparisons test. Control vs. *Prdm16* cKO, p=0.0041. *Prdm16 cKO* vs. *Prdm16-Vcam1* DKO, p=0.0055.

Next, to examine whether Prdm16 inhibition of Vcam1 is involved in the postnatal disappearance of radial glia and the ending of embryonic cortical neurogenesis, we generated *Prdm16-Vcam1* double cKO (*hGFAP-Cre^Tg/+^; Prdm16^fl/fl^, Vcam1^fl/fl^*, DKO) mice. Interestingly, the radial process persistence phenotype is partially rescued in DKO mice (Fig. 6B, E), suggesting that Prdm16 promotes the retraction of radial glia basal processes by inhibiting Vcam1. We next examined wholemount en-face preparations of the V-SVZ of one-month-old DKO mice to assess the abundance of NSCs. We found that the densities of NSCs and ependymal cells were restored to control levels in *Prdm16-Vcam1* DKO mice (Fig. 6C, F).

We further examined whether Prdm16 repression of Vcam1 is involved in the ending of cortical neurogenesis. Interestingly, we found that the postnatal persistence of cortical neurogenesis phenotype observed in *Prdm16* cKO mice was completely rescued in *Prdm16-Vcam1* DKO mice (Fig. 6G-H), suggesting that the elevation of Vcam1 levels in *Prdm16* cKO mice is essential for the postnatal continuation of cortical neurogenesis.

Given the Vcam1 rescue results (Fig. 6), and the potential role of Prdm16 as an epigenetic regulator that regulates gene expression, we examined whether Prdm16 regulates *Vcam1* transcription by scRNAseq. Surprisingly, *Vcam1* mRNA levels do not differ between *Prdm16* cKO and control mice (Supplementary Fig. 7) but Vcam1 protein levels are elevated in *Prdm16* cKO mice (Fig. 6 A, D), suggesting that Prdm16 does not directly regulate *Vcam1* transcription.

Together, we propose the following working model based on our results (Fig. 7): Vcam1 levels are high in embryonic NSCs/radial glia. Prdm16 induces a reduction in Vcam1 level postnatally, triggering the disappearance of radial glia and the ending of cortical neurogenesis. In *Prdm16* cKO mice, elevated Vcam1 levels drive the persistence of radial glia in adulthood and the continuation of cortical neurogenesis for a longer period postnatally. In *Prdm16-Vcam1* DKO mice, Vcam1 is absent, and the transition can occur. Vcam1 levels, regulated by Prdm16, thus determine whether radial glia stay in the embryonic NSC stage or progress to the adult NSC stage and whether embryonic neurogenesis continues or ends. These results fill a key knowledge gap in NSC biology and may inform future studies directed toward improving NSC-based therapies for neurological disorders.

**Figure 7.**
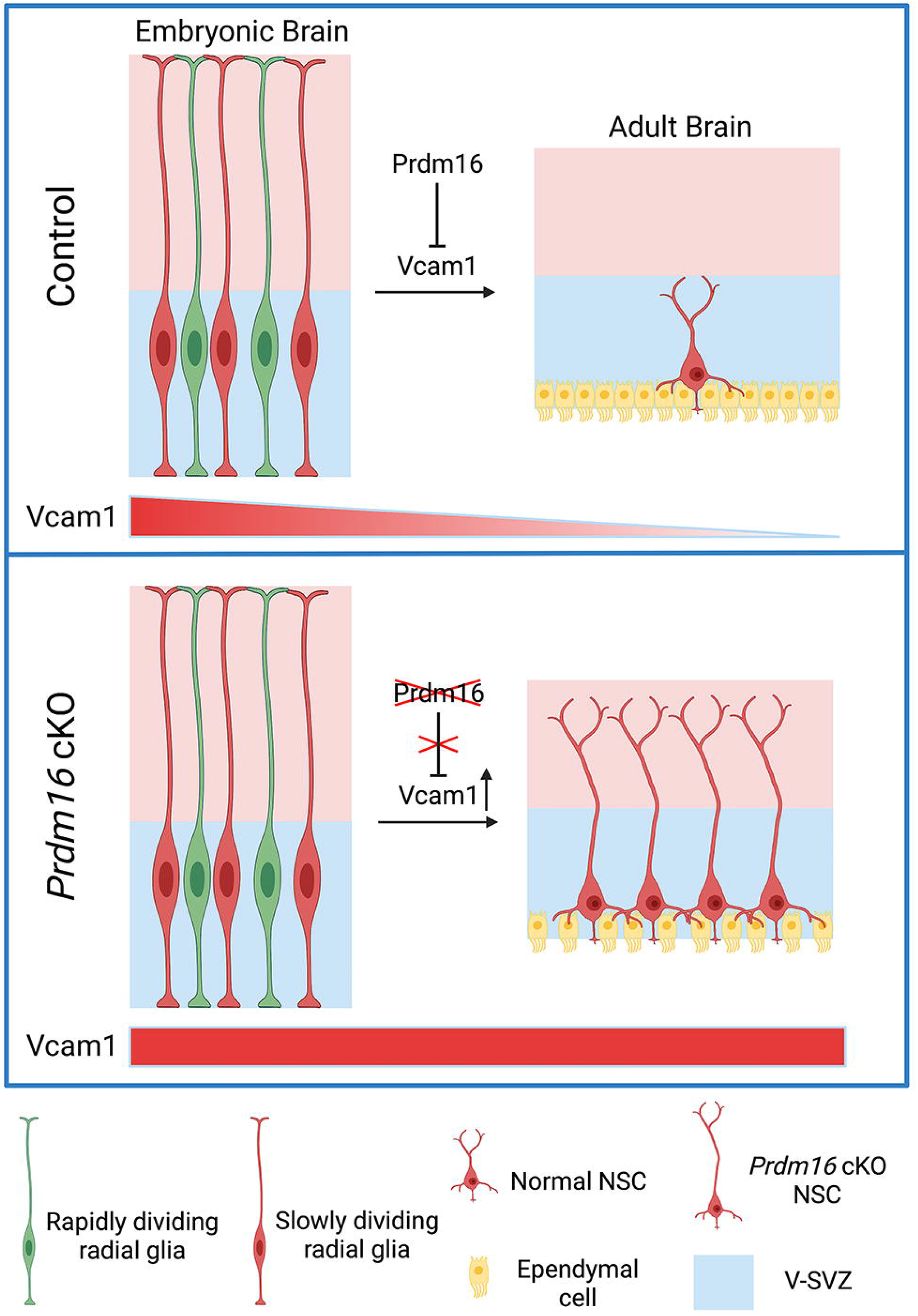
A working model of the roles of Prdm16 and Vcam1 in the transition from the embryonic to the adult NSCs. Embryonic slowly dividing radial glia become the adult type B cells postnatally. Vcam1 is highly expressed in the embryonic V-SVZ. In control mice, Prdm16 induces a reduction in Vcam1 levels postnatally, triggering the transition from embryonic radial glia to adult NSCs. In Prdm16 cKO mice, the expression of Vcam1 stays high postnatally, leading to the persistence of radial glia and the continuation of cortical neurogenesis.

## Discussion

In this study, we identified novel roles for Prdm16 and Vcam1 in promoting the postnatal disappearance of radial glia and the ending of cortical neurogenesis. These results may offer possible explanations for the differences in neurogenesis among vertebrate species. For example, fish display more prominent and widespread adult neurogenesis than mammals^62^, and the changes in evolution that led to this difference have not been identified. Interestingly, in zebrafish, *Prdm16* is not expressed after egg hatching^63^, unlike in mice, where *Prdm16* continues to be expressed by adult NSCs. In zebrafish, radial glia with basal processes persist in adulthood^64^, similar to *Prdm16* cKO mice. These observations are consistent with an evolutionarily conserved role of Prdm16 in the postnatal retraction of radial glia. Perhaps the postnatal expression of *Prdm16* first occurred in a species that appeared later than fish and earlier than mammals, leading to the retraction of the radial processes of embryonic NSC/radial glia, precluding postnatal cortical neurogenesis and the radial migration of new cortical neurons. Thus, adult neurogenesis is more limited in mammals than in fish.

ScRNAseq provides insight into the heterogeneity of cell populations. Both embryonic and adult NSCs have been extensively studied with scRNAseq^48–51,65–73^. However, the heterogeneity of NSCs during the transition from the embryonic to the adult stage is less clear. A pioneering study established the first single-cell transcriptomes of the transitional V-SVZ^48^. However, this effort profiled 2,143 cells from one-month-old V-SVZ, including only dozens to hundreds of cells from each major cell type. Here, we reported the transcriptomes of 43,306 one-month-old V-SVZ cells, providing high coverage of all major cell types. This comprehensive dataset allows for gene expression analyses of subtypes of cells in the V-SVZ, including diverse populations of activated and quiescent NSCs. This resource will be valuable for future studies exploring cell population changes during the transition from the embryonic to the adult V-SVZ.

In summary, we identified Prdm16 and Vcam1 as novel molecules that promote the postnatal disappearance of radial glia and the ending of cortical neurogenesis. Our study sheds new light on a poorly understood stage of NSC development and may contribute to the improvement of NSC-based therapies in the future.

## Acknowledgments

We thank Dr. Patrick Seale for sharing the Prdm16 antibody, Drs. Bruce Spiegelman and Paul Cohen for sharing the Prdm16 flox mice, Drs. Fiona Doetsch, Daniel Levy, Lindsay De Biase, Michael Sofroniew, Harley Kornblum, and Aparna Bhaduri for advice, Amy Gleichman and S. Tom Carmichael for assistance with imaging, Ranmal Samarasinghe for antibodies, and Brenda Urias for assistance with the illustrations. This work is supported by the NIH/NIMH T32MH073526 and the Achievement Rewards for College Scientists Foundation Los Angeles Founder Chapter to M. I. G., the UCLA Eli and Edythe Broad Center of Regenerative Medicine and Stem Cell Research (BSCRC) postdoctoral training grant to J.L., the NIH/NINDS NS111378, NS117148 and the NIH/NICHD HD100298 to X.Y., the NIH/NINDS R00NS089780, R01NS109025, the NIH/NIA R03AG065772, NIH/NICHD P50HD103557, National Center for Advancing Translational Science UCLA CTSI Grant UL1TR001881, the BSCRC Innovation Award, the Friends of the Semel Institute for Neuroscience & Human Behavior Friends Scholar Award, the W.M. Keck Foundation Junior Faculty Award, UCLA Jonsson Comprehensive Cancer Center and BSCRC Ablon Scholars Award to Y. Z.

## Author Contributions

J. L., M.I.G., and Y. Z. conceived the project, performed the experiments, and analyzed the data. A.J.Z. managed the mouse colony and performed some immunohistochemistry experiments. G.D., I.S.A., and X.Y. contributed to scRNAseq experiments and data analysis. B.G.N. contributed to the experimental design and provided reagents. A C-S and A A-B assisted with the whole-mount experiments. A A-B contributed to the experimental design. J.L., M.I.G., and Y.Z. wrote the paper. All authors read the manuscript.

## Methods

### Experimental animals

All experimental animal procedures were approved by the Chancellor’s Animal Research Committee at the University of California, Los Angeles and conducted in compliance with national and state laws and policies. We used mice group-housed in standard cages (2-3 adult mice, or 1-2 adults with a litter of pups per cage). Rooms were maintained on a 12-hour light/dark cycle. Euthanasia was performed during the light cycle. We used the following mouse strains; *hGFAP-Cre* (JAX004600), *Prdm16-* floxed (Spiegelman lab), and *Vcam1*-floxed (JAX007665). We crossed the strains to obtain control: *Prdm16^fl/fl^; Prdm16* single knockout: *hGFAP-Cre^+/-^; Prdm16^fl/fl^;* and *Prdm16-Vcam1* double knockout mice: *hGFAP-Cre^+/-^; Prdm16^fl/fl^; Vcam1^fl/fl^*. We also examined *Prdm16* heterozygous mice (*hGFAP-Cre^+/-^; Prdm16^fl/+^*) and did not observe any defect. We used both male and female mice in approximately equal proportions in the experiments and recorded data from male and female mice separately. We did not observe any sex-dependent effect in any assays and therefore combined data from both sexes.

### Immunohistochemistry on coronal brain sections

Mice were anesthetized with isoflurane and transcardially perfused with saline followed by 4% PFA. Brains were removed and further fixed in 4% PFA at 4°C overnight. The brains were washed with PBS and cryoprotected in 30% sucrose at 4°C for two days before being immersed in OCT (Fisher, cat#23-730-571) and stored at −80°C. Brains were sectioned on a cryostat (Leica) and 30μm floating coronal sections were blocked and permeabilized in 10% donkey serum with 0.2% Triton X-100 in PBS and then stained with primary antibodies against PRDM16 (Patrick Seale lab, dilution 1:100), PRDM16 (R&D system, AF6295-SP, 1:200), GFAP (Biolegend, 829401, 1:1500 and Dako, Z0334, 1:1500), Nestin (Aveslabs, NES-0020, 1:500), Vcam1 (BD Biosciences, 550547,1:100), Dcx (Abcam, ab18723, 1:500), BrdU (Abcam, ab6326, 1:500), Parvalbumin (Novus Biologicals, NB120-11427SS, 1:500), TH (Sigma, MAB318, 1:400), and Calretinin (Sigma, ZMS1073-25UL, 1:400) at 4°C overnight. Sections were washed three times with PBS and incubated for 2 hours at room temperature with secondary antibodies followed by three additional PBS washes. The sections were then mounted on Superfrost Plus micro slides (Fisher, cat#12-550-15) and covered with mounting medium (Fisher, cat#H1400NB) and glass coverslips. The coronal brain sections were imaged with a Zeiss widefield fluorescence microscope or a Zeiss LSM800 confocal microscope with 20, 40, and 63x lenses.

### Immunohistochemistry on the SVZ whole-mount preparations

We prepared the V-SVZ whole-mount preparations and performed immunostaining according to published protocols^47^. Briefly, we euthanized the mice and dissected out the brains. We then carefully expose the lateral walls of the lateral ventricles and cut away other brain structures. We then fixed the whole-mounts, ventricle side up, with 4% PFA with 0.1% Triton-X100 overnight at 4°C. We then washed the whole-mounts 3 times with PBS with 0.1% Triton-X100, blocked in 10% normal donkey serum in PBS with 2% Triton-X100 at room temperature for 1 hour, and stained with primary antibodies against Vcam1 (BD Biosciences, 550547, 1:100), γ-Tubulin (Sigma, T5192-25UL, 1:1000), and SOX2 (R&D Systems, AF2018, 1:1000) for 48 hours at 4°C. We then quickly rinsed the whole-mounts with PBS three times, stained them with fluorescent secondary antibodies for 48 hours at 4°C, and quickly rinsed them with PBS three times. After staining, we sub-dissected the whole-mounts to preserve only the lateral wall of the lateral ventricle as a sliver of tissue 200-300 μm thick and mounted the tissue in FluorSave^TM^ Reagent mounting medium on glass slides with glass coverslips. We imaged the whole-mounts with a Zeiss LSM800 confocal microscope with a 40x lens.

### BrdU pulse chase

We intraperitoneally injected 50 mg/kg BrdU into pregnant female mice daily for three consecutive days at E15.5-17.5. The progeny was sacrificed by transcardial perfusion at P21. The brains were processed for immunohistochemistry as described above with the following modifications: the brain sections were first treated with 2M HCl for 30 min at 37°C followed by treatment with 0.1M boric acid for 10 min before adding the primary antibodies.

### Sample and library preparation for scRNAseq

The V-SVZ of the lateral walls of the lateral ventricles from one-month-old (P31-33) mice were dissected according to published protocols^47^. The V-SVZs from 2-3 mice including both males and females were combined as a biological replicate. Single-cell suspensions were prepared as described in^74^. Briefly, a 2 mg/ml papain solution was pre-activated at 37°C for 20 min. The V-SVZ samples were then digested in the papain solution at 30 °C for 30 min with a shaking speed of 60 rpm in a shaker. The tissue sample is then mechanically dissociated by trituration with a Pasteur pipette 10~15 times per round for three rounds, washed, and centrifuged with an OptiPrep density gradient to remove myelin and cellular debris. After centrifugation, the dense white layer and all layers above were carefully aspirated. The lower layers were pelleted at 200g for 3min, resuspended in 1 ml PBS with 0.04% bovine serum albumin (0.04% BSA-PBS), filtered through 40 μm strainers, and resuspended in 0.04% BSA-PBS at 700-1,200 cells/μl for single-cell capture. Cell viability was assessed by Trypan Blue staining and a final cell count was performed before single cell capture. The single-cell suspensions with > 60% viable cells were used for single-cell capture. The 10x Genomics Chromium Next GEM Single Cell 3’ Reagent Kits v3.1 were used for GEM generation, barcoding, and library preparation according to the manufacturer’s instructions. Reagents were prepared to aim to capture 7,000-10,000 cells per sample. The TapeStation was used for cDNA and library quality control. The libraries were sequenced with an Illumina NovaSeq sequencer with S2 x100 cycles at the UCLA Technology Center for Genomics and Bioinformatics to obtain 90,871 reads per cell on average.

### scRNAseq data analysis

Reads were mapped to the mouse genome (GRCm38) with cellranger count (version 6.1.2). Feature-count matrices generated by Cell Ranger were next processed with Seurat version 4.1.0^75^ for filtering, clustering, and differential gene expression analysis. Low-quality cells with fewer than 500 features or higher than 10% mitochondrial genes were filtered out. Potential doublets with more than 7000 features were also removed. In total, we obtained transcriptome data for 36,420 V-SVZ cells that passed quality control, including 21,939, 14,481, and 6,886 cells from control, *Prdm16* cKO, and *Prdm16-Vcam1* DKO mice, respectively. The data were then normalized and scaled using the “LogNormlize” method in the Seurat package. We then found the top 2000 highly variable features and used the top 30 principal components for dimensional reduction. We used the FindIntegrationAnchors function in Seurat to anchor and integrate individual samples together and removed batch effects. We next clustered the cells using 0.15 resolution and used UMAP for dimension reduction. We assigned cell type identity to clusters based on known cell-type specific marker genes^48,50,65,67,76^ and identified differentially expressed genes using the FindMarkers function in Seurat. To calculate pseudobulk gene expression for each cell type, we used the AverageExpression function in Seurat. For the analysis of the E14.5 dataset^48^, we first integrated data from all E14.5 samples using the IntegrateData function in Seurat. We then removed low-quality cells with fewer than 500 features or higher than 6% mitochondrial genes, and potential doublets with more than 6000 features. Next, we identified the NSC cluster based on the expression of *Nestin, Fabp7*, and *Vcam1*. For GO and protein-protein interaction network analysis, we used string-db.org with default settings^77^.

### Gene Set Enrichment Analysis (GSEA)

To confirm the GO analysis results, we also performed Gene Set Enrichment Analysis. We downloaded GSEA (version 4.2.3) software from www.gsea-msigdb.org. We used the default settings with the following exceptions: we first ranked the *Prdm16* cKO NSC *vs*. control NSC differentially expressed genes by fold change, then ran GSEAPreranked with this ranked gene list. We used c5.go.bp.v7.5.1.symbols.gmt as “Gene sets database”. We obtained similar results by GSEA as the GO term enrichment analysis.

To compare *Prdm16* cKO NSC *vs*. control NSC differentially expressed genes and slowly dividing radial glia *vs*. adult NSC differentially expressed genes by GSEA, we built our own gene sets of slowly dividing radial glia *vs*. adult NSC differentially expressed genes. We generated these gene sets by comparing the Borrett et al. E14.5 *vs*. P20-61 NSC cluster data. We first merged the differentially expressed genes from *Prdm16* cKO NSC *vs*. control NSC and radial glia *vs*. adult NSCs. Next, we made a radial glia-enriched gene set and an adult NSC-enriched gene set from the merged radial glia vs. adult NSC differentially expressed genes. We ran GSEAPreranked with the ranked *Prdm16* cKO NSC *vs*. control NSC differentially expressed genes to assess enrichment of genes from the radial glia-enriched gene set and adult NSC-enriched gene set.

### Quantification and statistical analysis

Image analyses were performed with the experimenter blinded to the genotype of the mice with the following exception: due to the enlarged ventricle phenotype of the Prdm16-knockout mice, blinding of images including the ventricles was not possible. The numbers of mice and replicates are described in the figure legends. scRNAseq data were analyzed as described above. All non-RNA-seq data analyses were conducted using R and the Prism 8 software (Graphpad). The normality of data was tested by the Shapiro-Wilk test. For data with normal distribution, Welch’s t-test was used for two-group comparisons and one-way ANOVA with Tukey’s multiple comparison test was used for multi-group comparisons. For data that deviate from a normal distribution, the Wilcoxon signed-rank test (for paired data) was used. Data from technical replicates from the same mouse were averaged and used as a single biological replicate in statistical tests. An estimate of variation in each group is indicated by the standard error of the mean (S.E.M.). * p<0.05, ** p<0.01, *** p<0.001.

### Data and code availability

We will deposit all RNA-seq data to the Gene Expression Omnibus. This study does not generate new codes.

