## Supplementary figures and images for "Prdm16 and Vcam1 regulate the postnatal disappearance of embryonic radial glia and the ending of cortical neurogenesis"

### Supplementary Figure 1

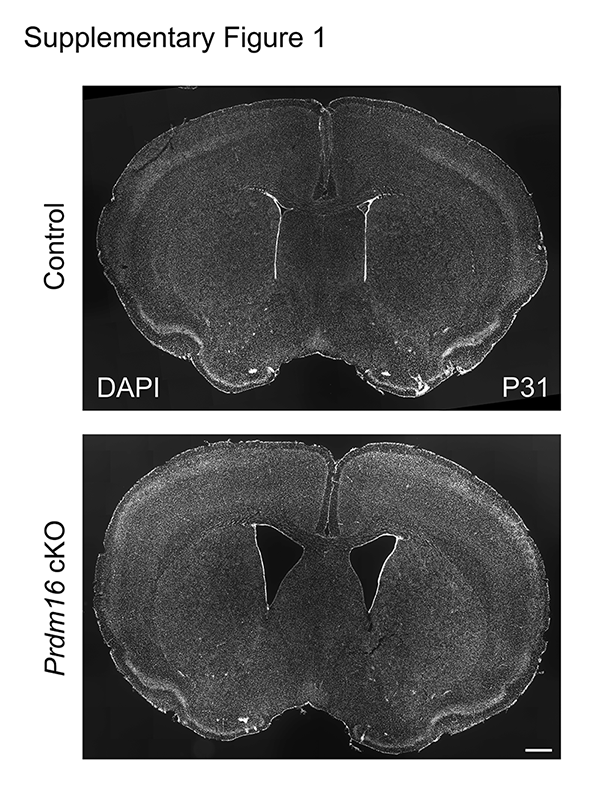

### Supplementary Figure 2

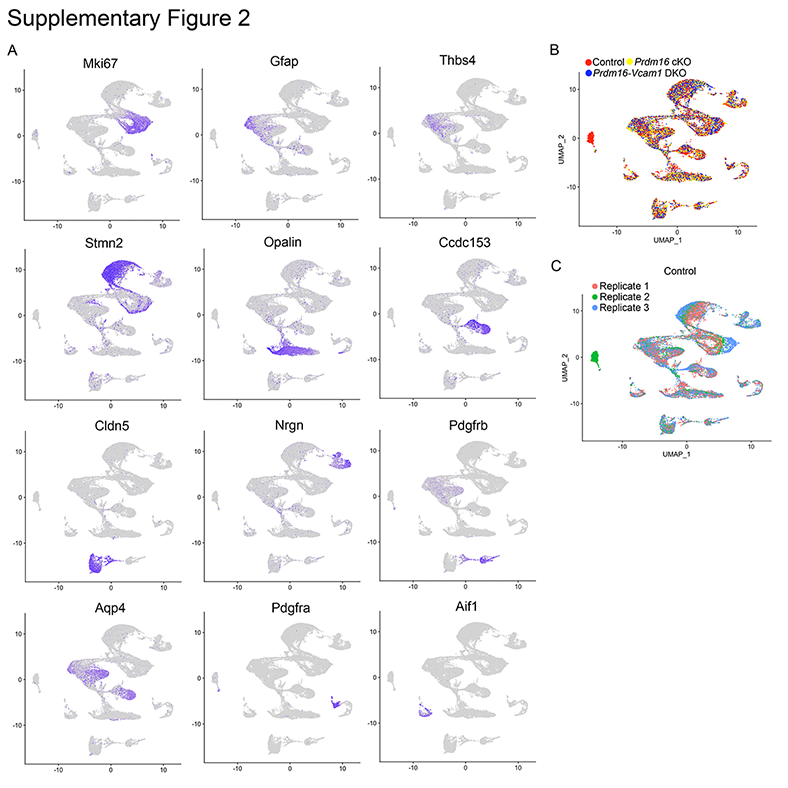

### Supplementary Figure 3

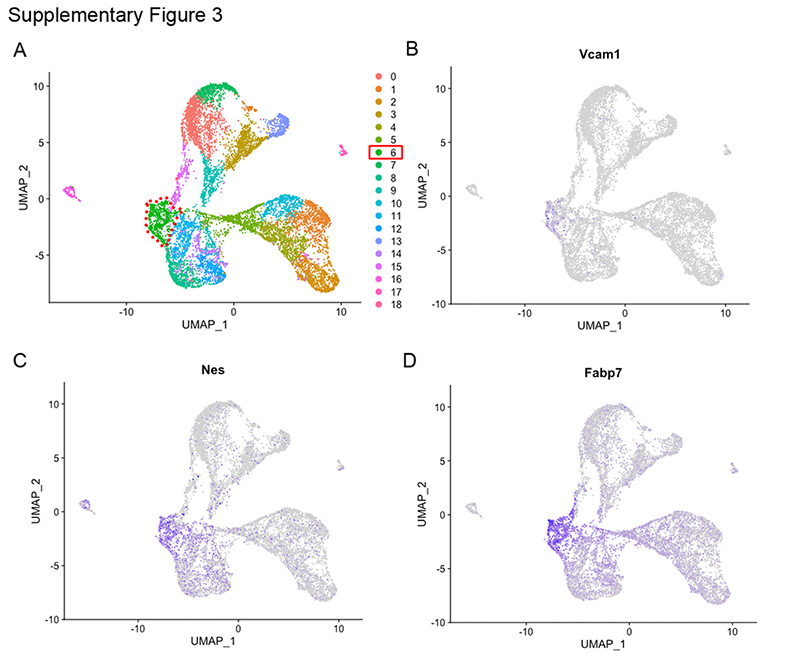

### Supplementary Figure 4

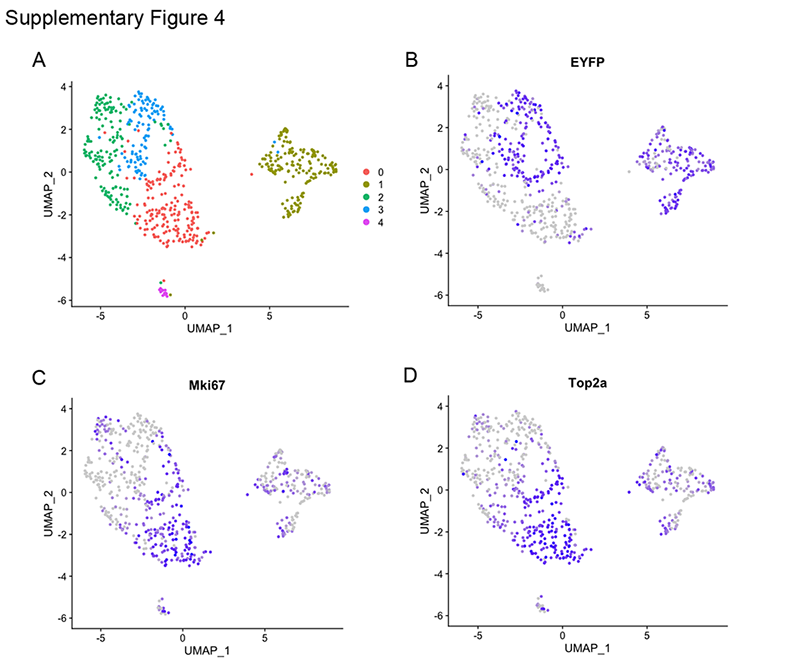

### Supplementary Figure 5

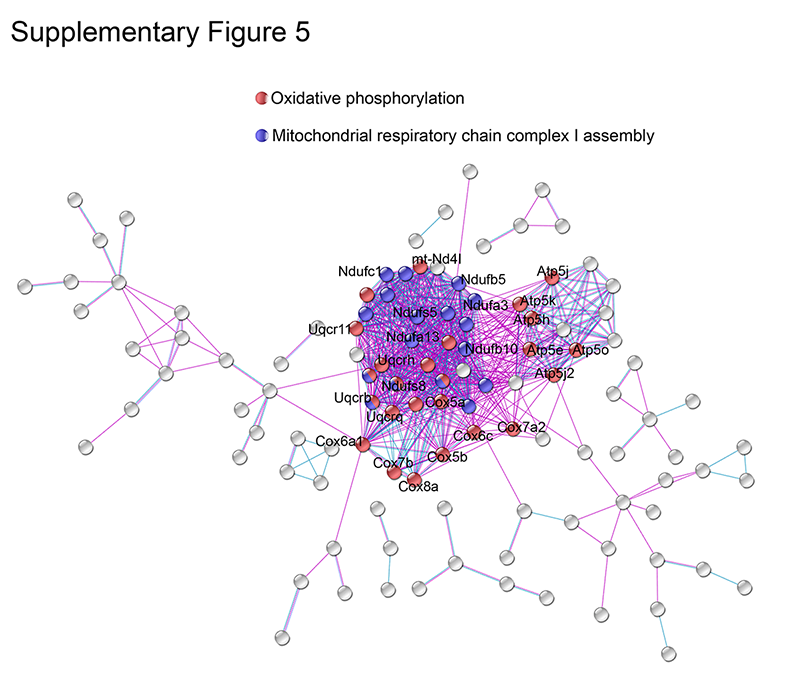

### Supplementary Figure 6

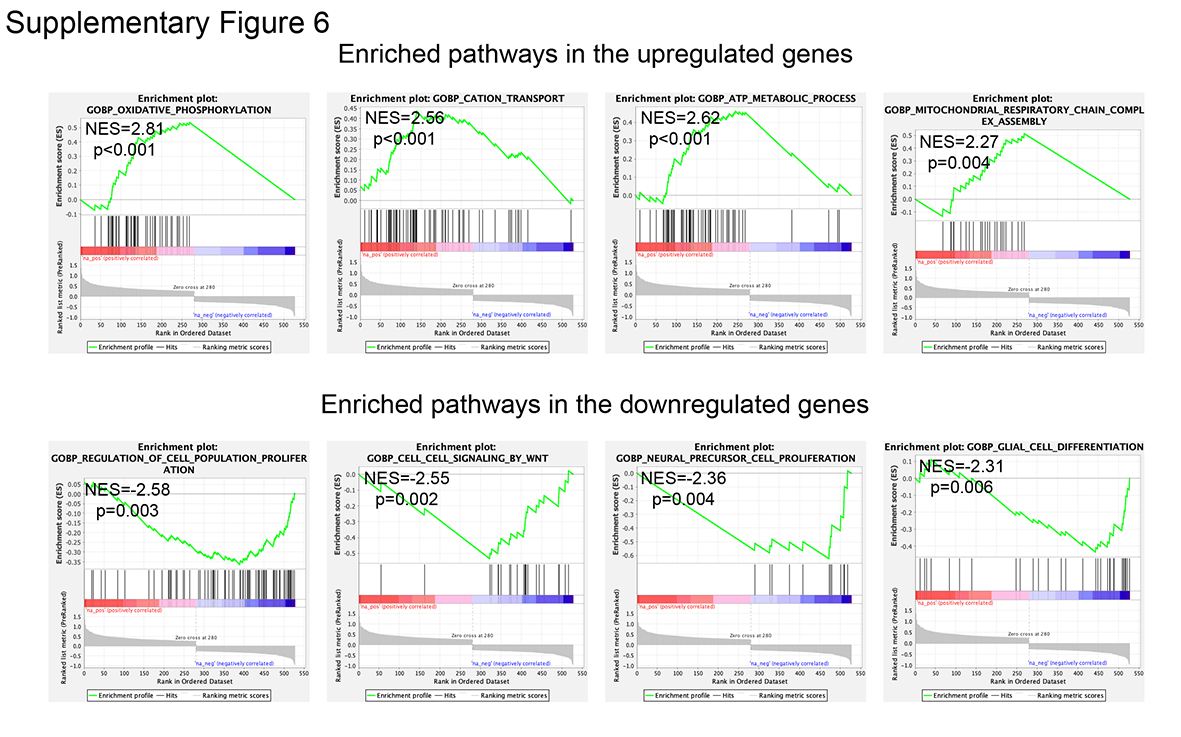

### Supplementary Figure 7

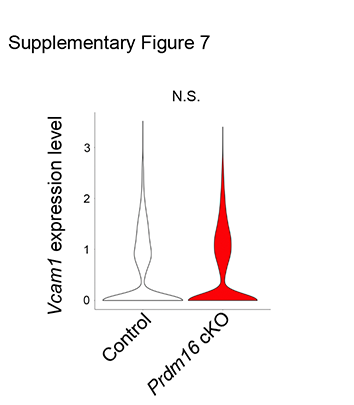
