## Supplementary Information for "Prdm16 and Vcam1 regulate the postnatal disappearance of embryonic radial glia and the ending of cortical neurogenesis"

**Supplementary Table 1. Pseudobulk gene expression levels in each ventricular-subventricular zone cell type in one-month-old *Prdm16* cKO, *Prdm16-VCAM1* DKO, and control mice.**

Gene expression is shown as transcripts per million.

**Supplementary Table 2. Genes differentially expressed in each cell type in *Prdm16* cKO vs control mice**

P_val, nominal P-value. P_val_adj, P-values adjusted for multiple comparisons. Ave_log2FC, log2 fold change. Pct. 1, percentage of cells expressing the gene in Prdm16 cKO. Pct.2, percentage of cells expressing the gene in control.

**Supplementary Table 3. Enriched gene ontology terms in genes up- and down-regulated in NSCs in *Prdm16* cKO compared with control mice**

**Supplementary Figure 1. The hydrocephalus phenotype in *Prdm16* cKO mice**

Scale bar: 500 μm

**Supplementary Figure 2. Expression of cell-type markers by major cell clusters in the scRNAseq dataset**

(A) We used the expression of the following cell-type markers to identify cell clusters: transient amplifying precursors (TAP, *Mki67*), neural stem cells (*Gfap, Thbs4*), neuroblasts (*Stmn2*), oligodendrocytes (*Opalin*), ependymal cells (*Ccdc153*), epithelial cells (*Cldn5*), neurons (*Nrgn*), pericytes (*Pdgfrb*), astrocytes (*Aqp4*), oligodendrocyte precursor cells (*Pdgfra*), microglia (*Aif1*). The expression of the marker genes on UMAP is shown.

(B, C) Control, *Prdm16* cKO, and *Prdm16-Vcam1* DKO samples contain cells from all clusters except for cluster 6 (red in B), which is only detected in one of the three control samples. We excluded cluster 6 from further analyses.

**Supplementary Figure 3. Identification of the radial glia cluster in the E14.5 scRNAseq dataset**

Cluster 6 (outlined in red) in (A) expresses *Vcam1* (B), *Nestin* (C), and *Fabp7* (D) and was identified as the radial glia cluster.

**Supplementary Figure 4. Subclusters of radial glia in the E14.5 scRNAseq dataset**

(A) UMAP showing 5 subclusters of radial glia in the E14.5 scRNAseq dataset.
(B) Expression of Emx1-EYFP suggests that subclusters 1 and 3 are dorsal radial glia and subclusters 0 and 2 are lateral radial glia

(C, D) Expression of the proliferative cell markers Mki67 and Top2a suggests that subcluster 2 is pre-B radial glia and subcluster 0 is non-pre-B radial glia.

**Supplementary Figure 5. Protein-protein interaction network analysis**

Protein-Protein interaction networks in the genes upregulated in the NSCs from *Prdm16* cKO mice. The highlighted GO terms are oxidative phosphorylation (red) and mitochondrial respiratory chain complex I assembly (purple).

**Supplementary Figure 6. Gene set enrichment analysis of differentially expressed genes between *Prdm16* cKO and control NSCs**

**Supplementary Figure 7. *Vcam1* mRNA expression level in the NSCs of one-month-old *Prdm16* cKO and control mice as determined by scRNAseq**
